# Investigation of lanthanide-dependent methylotrophy uncovers complementary roles for alcohol dehydrogenase enzymes

**DOI:** 10.1101/329011

**Authors:** Nathan M. Good, Olivia N. Walser, Riley S. Moore, Carly J. Suriano, Anna F. Huff, N. Cecilia Martínez-Gómez

**Affiliations:** Department of Microbiology and Molecular Genetics, Michigan State University

## Abstract

The discovery that methylotrophic bacteria can utilize lanthanides as catalysts for methanol metabolism has opened new areas of biology and biochemistry. Recent studies of lanthanide-dependent enzymes have focused on biochemical and kinetic properties or the regulation of encoding genes. Kinetic analysis of a pyrroloquinoline quinone methanol dehydrogenase, XoxF1 (MexAM1_1746), from the model methylotroph *Methylobacterium extorquens* AM1 confirms the use of different lanthanides as cofactors and formaldehyde as a product of methanol oxidation, showing that not all XoxF MDH produce formate as the only end product *in vivo*. The dephosphotetrahydromethanopterin pathway for formaldehyde oxidation is still required for lanthanide-methylotrophic growth, as a *fae* mutant does not grow with methanol in the presence of exogenous lanthanides. Increases of 15-22% in growth rate and 10-12.5% in growth yield are observed when *M. extorquens* AM1 is grown in the presence of lanthanides with methanol. RNA-sequencing transcriptomics indicates remodeling of methanol, formaldehyde and formate oxidation gene expression, and targeted metabolomics shows increased accumulation of intracellular formate and decreased pools of several assimilatory intermediates. Methanol sensitivity growth assays show that the lanthanide-dependent pyrroloquinoline quinone alcohol dehydrogenase ExaF (MexAM1_1139), but not XoxF1, can reduce formaldehyde toxicity when lanthanides are present, providing evidence of a role for ExaF during lanthanide-dependent methylotrophy. We conclude from these results that lanthanide-dependent methylotrophy is more efficient than calcium-dependent methylotrophy in *M. extorquens* AM1, and that this change is due, at least in part, to the lanthanide-dependent enzymes XoxF1 and ExaF.

**IMPORTANCE:** Lanthanides serve as cofactors for pyrroloquinoline quinone containing alcohol dehydrogenase enzymes in methylotrophic bacteria. The present study addresses a fundamental gap in our understanding of how lanthanides impact metabolism, including a detailed assessment of the metabolic modifications to accommodate enhanced efficiency during methylotrophy. Kinetic characterization of XoxF1 provides a detailed description of the impact of diverse lanthanides on catalytic function for a lanthanide-dependent methanol dehydrogenase. We further show that the lanthanide-dependent ethanol dehydrogenase ExaF can oxidize formaldehyde *in vivo*, revealing complementary roles for these enzymes. This study provides novel insight into the effects of lanthanides on bacterial metabolism, highlighting the implementation of multiple, redundant and complementary oxidation systems.

## INTRODUCTION

A direct link between the lanthanide elements (Ln) and microbial metabolism has been firmly established with the discovery of pyrroloquinoline quinone (PQQ)-dependent alcohol dehydrogenases (ADH) that contain a Ln atom in the active site in methylotrophic bacteria (1– 5). Thus far, Ln-ADH can be grouped by their phylogeny and primary substrate as either XoxF-type methanol dehydrogenases (MDH) or ExaF-type ethanol dehydrogenases (EtDH). Reports have provided evidence for XoxF having roles as both a periplasmic methanol and formaldehyde-oxidation system (6) and as a regulator of expression of the calcium (Ca) dependent MxaFI MDH (6, 7). MxaFI MDH has been considered the canonical primary catalyst for methanol oxidation in Gram-negative methylotrophs (8, 9). MxaFI is a heterotetramer that contains PQQ as a prosthetic group and coordinates a calcium ion (10–12). The discovery that Ln are incorporated into the active site of XoxF MDH in place of Ca, allowing catalytic function, has prompted the reexamination of methanol oxidation in methylotrophic bacteria (3, 13, 14). In addition, we reported the first Ln-dependent EtDH ExaF, showing that Ln can impact multi-carbon as well as one-carbon metabolism (5). This finding was later corroborated by the discovery of differentially regulated Ca-and Ln-dependent EtDHs in *P. putida* KT2440 (15). A growing number of studies indicate a wide-spread role for Ln in microbial metabolism that we are only beginning to understand.

To date, only a few XoxF MDH have been kinetically characterized (1, 3, 4, 16). The only reported crystal structure of XoxF MDH is from the methanotrophic Verrucomicrobium *Methylacidiphilum fumariolicum* SolV (3). Phylogenetic analyses show there are at least five distinct families of XoxF MDH (13), and while it has been suggested that all XoxF MDH may exhibit similar kinetic properties, reported data for these enzymes are currently inadequate to support such a broad characterization. Catalytic properties observed for XoxF MDH from *M. fumariolicum* SolV and ExaF EtDH from *M. extorquens* AM1 reveal higher catalytic efficiencies compared to homologous Ca-containing enzymes (13) and the capability to produce formate as the end product of methanol oxidation *in vivo* (3, 6). The genome of *M. extorquens* AM1 contains two *xoxF* genes, named *xoxF1* and *xoxF2*, respectively (17). Either gene product is capable of supporting methanol growth with exogenous Ln, but XoxF1 is the primary methanol oxidation system for Ln-dependent methylotrophy (18). Increased catalytic function was observed for XoxF1 purified with lanthanum (La^3+^) as a cofactor, leading to the description of the enzyme as La^3+^-dependent (2). The only detailed kinetic analysis available for XoxF1, however, was conducted with enzyme from culture grown in the absence of Ln (7). I*n vivo* evidence is suggestive of XoxF1-catalyzed formaldehyde oxidation in starving cells fed methanol. However, metabolically active cells were not used in those studies and, nor was the regulatory role of XoxF1 taken into account (6, 19). Since these studies were done in the absence of Ln, it is currently unknown how the catalytic properties of XoxF1 could change *in vivo*. Due to the relevance of this enzyme for Ln-dependent methylotrophy, and the lack of fundamental information available for Ln-dependent enzymes, particularly with other Ln cofactors, a detailed kinetic study of XoxF1 MDH is needed.

Defining the kinetic differences of Ln-dependent MDH, and assessing the impacts they have on downstream metabolism, is an important first step to better understanding the implications of these metals in biology. Methanol growth in *M. extorquens* AM1 requires the dephosphotetrahydromethanopterin (H_4_MPT) pathway to link highly reactive formaldehyde produced from methanol oxidation to the pterin carbon carrier (20). This pathway is indispensable for growth on methanol, and functions for both formaldehyde oxidation and NAD(P)H and formate generation (21, 22). If XoxF1 exhibits catalytic properties similar to the enzyme from *M. fumariolicum* SolV and ExaF, is the H_4_MPT pathway dispensable? If, instead, XoxF produces formaldehyde from methanol oxidation, what then, if any, is the role of ExaF during Ln-dependent methylotrophy? The potential catalytic efficiency of Ln-ADH raises the possibility of wide-spread changes to methylotrophic metabolism.

Global studies of gene expression and metabolite profiles would provide valuable insight into the impacts of Ln on methylotrophy beyond the regulatory change from a Ca-MDH to a Ln-MDH, referred to as the “lanthanide switch” (23). A recent study utilizing gene expression studies comparing methylotrophy with and without La^3+^ for *Methylobacterium aquaticum* strain 22A reported down-regulation of formaldehyde oxidation and PQQ synthesis genes (19). This same study described a higher formaldehyde oxidation rate in resting cells when grown in the presence of La^3+^, again indicative of a Ln-dependent change to formaldehyde metabolism. Detailed characterization of the metabolomics profile, coupled with gene expression studies of actively growing cells, would provide further insight into the metabolic effects of Ln on methylotrophy.

In this study, we biochemically characterize XoxF1 MDH from the model methylotroph *M. extorquens* AM1, which utilizes the serine cycle and ethylmalonyl-CoA pathway (EMC) for carbon assimilation (14). We confirm that XoxF1 MDH from *M. extorquens* AM1 is dependent on Ln metals for catalytic function, that formaldehyde is the product of methanol oxidation, and that it exhibits properties similar to those reported for MxaFI enzymes. In support of this, we show that the H_4_MPT pathway is required during Ln methylotrophy. RNA-seq transcriptomics and targeted metabolomics analyses indicate marked modifications to methanol and formate oxidation, as well as significant alterations to assimilatory pathways. Together, these modifications to methylotrophy result in a faster growth rate and higher growth yield compared to Ln-independent growth with methanol. Finally, we show that ExaF EtDH reduces methanol sensitivity in a *fae* deletion mutant strain by alleviating toxic accumulation of formaldehyde, confirming a contributing role in formaldehyde oxidation during Ln-dependent methylotrophy.

## RESULTS

### Enzyme kinetics of XoxF1 MDH - a *bona fide* Ln-dependent methanol dehydrogenase

XoxF1 from *M. extorquens* AM1 is a functional La^3+^-dependent MDH and the primary oxidation system utilized for Ln-dependent growth with methanol (2, 18). However, detailed kinetic parameters have only been reported for XoxF1 purified in the absence of La^3+^ (6). Further, catalytic properties of XoxF1 have not been studied with other Ln. The only other Ln-dependent MDH biochemically characterized thus far exhibits an incredible capacity to oxidize methanol, more than 10 times higher than any other reported PQQ MDH (3) and was shown to use lanthanides such as Ce^3+^ and Eu^3+^ (24). Interestingly, this enzyme is a type II XoxF-MDH and only 51.9% identical to XoxF1 from *M. extorquens* AM1, a type V XoxF-MDH. Therefore, we kinetically characterized XoxF1 as an additional comparator to the few Ln ADHs reported thus far.

XoxF1 from *M. extorquens* AM1 was produced in cultures grown with methanol and 20 μM LaCl_3_. Using IMAC, histidine-tagged XoxF1 was enriched and purified to homogeneity after histidine tag cleavage via tobacco etch virus (TEV) protease (25, 26). Methanol oxidation was measured by monitoring the PMS-mediated reduction of DCPIP ((10, 18). Specific activity of cell-free extract containing XoxF1 with La^3+^ (XoxF1-La) was 67 nmol mg^−1^ min^−1^. PQQ was detected as a prosthetic group via UV-Visible spectroscopy, with a molar ratio of 1.2 mol PQQ per mol of enzyme [FIG 1D]. Metal content determined by ICP-MS showed that XoxF1-La contained La^3+^ in 1:1 molar ratio of metal to protomer.

**FIG 1.**
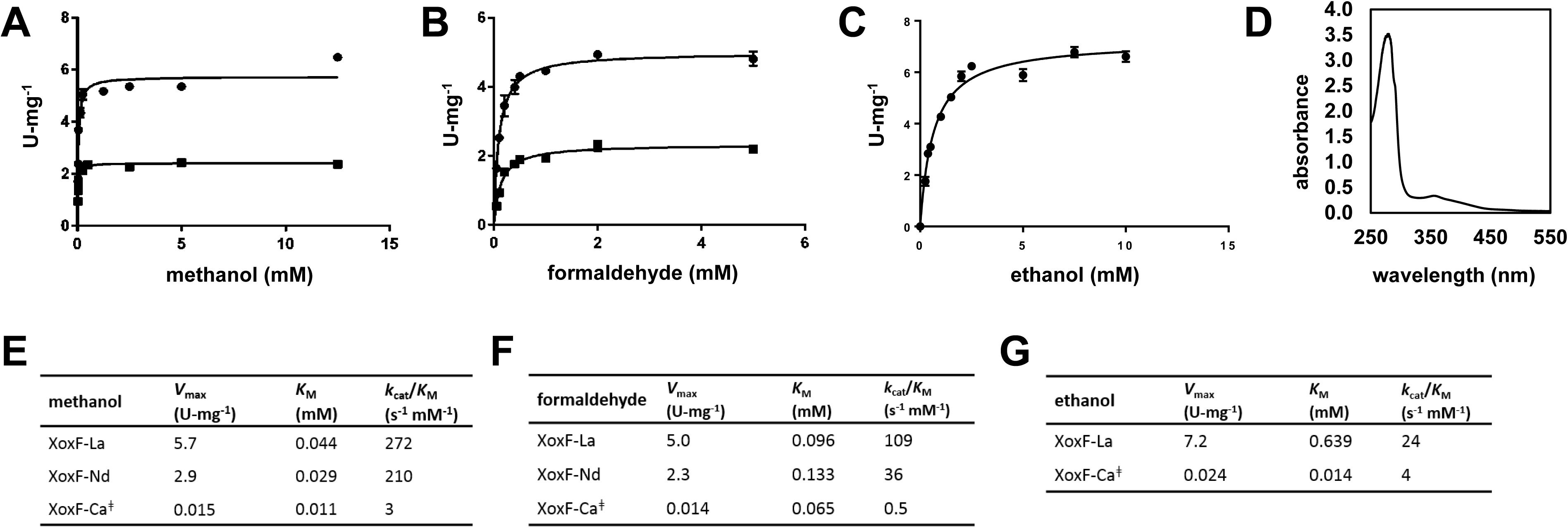
Biochemical characterization of Ln-dependent XoxF1 MDH. XoxF1 was purified and characterized with La^3+^ (circles) or Nd^3+^ (squares) as the metal cofactor. Michaelis-Menten plots for XoxF1 with methanol (A), formaldehyde (B), and ethanol (C) are shown. Values are the mean of triplicate measurements with error bars representing the standard deviation. PQQ was detected as the prosthetic group in XoxF1 by UV-Visible spectrum analysis (D), with a peak from 345-350 nm. Enzyme was prepared in 25 mM Tris-HCl, pH 8.0 at a concentration of 3.5 mg/ml. Kinetic parameters determined from plots A-C. XoxF-La, XoxF-Nd, XoxF-Ca are XoxF1 purified with lanthanum, neodymium, and without a lanthanide, respectively.

Kinetic parameters were measured for XoxF1-La [FIG 1] and compared to reported values for XoxF1 purified from cultures grown without Ln (6). With La^3+^ bound, the *V*_max_ of methanol oxidation via XoxF1 was 380-fold higher. The *K*_M_ for methanol was 4-fold higher. This resulted in a 91-fold increase in catalytic efficiency when La^3+^ is coordinated in the enzyme [FIG 1, A and E]. Increased enzyme function when La^3+^ was bound was observed for formaldehyde and ethanol oxidation as well [FIG 5 B-C, and F-G]. With formaldehyde as the substrate, the *V*_max_ increased 357-fold, the *K*_M_ increased less than 2-fold, and catalytic efficiency increased 218- fold. With ethanol as the substrate, *V*_max_, *K*_M_, and catalytic efficiency increased 300-fold, 45- fold, and 6-fold respectively. These results show a definitive increase in catalytic function of XoxF1 MDH when La^3+^ is available for incorporation into the active site, and indicate that oxidation rate, rather than increased substrate affinity, is cause of this increase.

Next, we purified XoxF1 from cultures grown with exogenous neodymium (Nd^3+^) and kinetically characterized the enzyme with methanol and formaldehyde as substrates [FIG 1 A-B, E-F]. ICP-MS metal determination of XoxF1-Nd protein resulted in a 0.5:1 lanthanide to protomer molar ratio. Reconstitution to a 1:1 Nd to protomer ratio was unsuccessful *in vitro*. Nd^3+^ content correlated with a ∼50% reduction in *V*_max_ compared to XoxF1-La with either methanol or formaldehyde as the substrate. Relative changes in *K*_M_ values were divergent and substrate-dependent. The *K*_M_ for methanol decreased 33%, while the *K*_M_ for formaldehyde increased 39%, indicating that Ln may differentially impact MDH affinity for a substrate.

Kinetic analysis shows that pure XoxF1 with Ln is capable of oxidizing formaldehyde *in vitro*, but it is not known if this activity is present *in vivo* as well. Such activity would imply production of formate via MDH in the periplasmic space when the cells are grown with methanol in the presence of Ln. Previous studies have relied on artificial or indirect methods to measure *in vivo* oxidation of methanol to formate. *Pol, et al.* compared substrate consumed to product formed in the dye-linked *in vitro* MDH assay using pure XoxF MDH from *M. fumariolicum* SolV. They reported a 2-fold increase in the amount of DCPIP reduced relative to methanol oxidized, indicating that the substrate was oxidized to formaldehyde and then to formate (3). An increased rate of formaldehyde accumulation was also observed in resting cells of *M. aquaticum* strain 22A when cultures were grown in the absence of La^3+^ than when in presence of La^3+^ (19). Based on these results, it has been proposed that all XoxF MDH may produce formate, and not formaldehyde, as the final oxidation product of methanol. Under methanol-limiting conditions, XoxF1-La or XoxF1-Nd reduced equimolar concentrations of DCPIP to methanol, indicating that formaldehyde, and not formate, is the final product of methanol oxidation for this enzyme *in vitro*.

### Formaldehyde-activating enzyme is required for Ln-dependent growth with methanol

Under Ln-free conditions, an *fae* mutant strain cannot grow with methanol as a carbon source due to the lack of enzymatic coupling of free formaldehyde to the carbon carrier H_4_MPT, resulting in reduced carbon flux to formate and toxic accumulation of formaldehyde (27, 28). If XoxF1 produces formate as the final oxidation product of methanol, Fae activity could be dispensable during growth with Ln. We tested the *fae* mutant strain for the ability to grow with methanol with and without La. No growth was observed for either condition with 15 mM or 125 mM methanol as the growth substrate [Fig 2A], indicating that Fae, and by association the H_4_MPT pathway, is still needed for Ln-dependent methanol growth. Carbon flow through the H_4_MPT pathway, therefore, is required for Ln-dependent methylotrophy providing *in vivo* evidence with a growing culture that XoxF1 catalyzes the production of formaldehyde, and not formate, from methanol.

**FIG 2.**
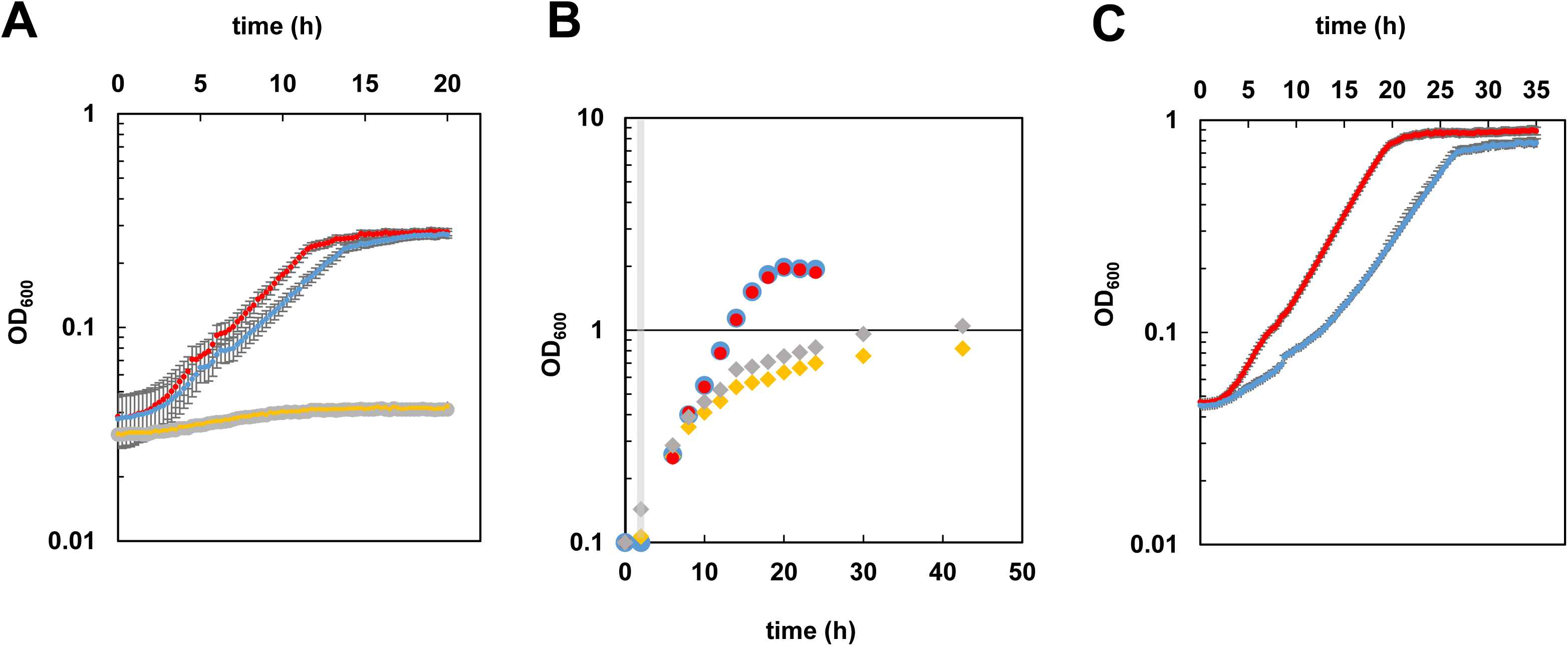
Growth of wild-type and *fae* mutant strains of *M. extorquens* AM1 in MP minimal medium with 15 mM (A) or 125 mM (C) methanol, in the presence [red, wild-type; gray, *fae*] or absence [blue, wild-type; yellow, *fae*] of 2 μM exogenous LaCl_3_. Cultures were grown in 48-well cell culture plates shaking at 548 rpm at 30°C in an EpochII plate reader. Optical density at 600 nm was measured in 15 minute intervals. Error bars represent SEM of four biological replicates. Not all error bars are visible due to the size of the marker representing the error. (B) Growth of wild-type and *fae* mutant strains on succinate with methanol addition after 2 hours of growth [gray bar]. All strains were grown in plastic 15 mL culture tubes. Growth was measured as the increase in culture absorbance at 600 nm. Data points are representative of three biological replicates with <5% variance. Wild-type [circles] and Δ*fae* [diamonds] mutant strains were grown in the presence [red, gray] or absence [blue, yellow] of 2 μM LaCl_3_.

An *fae* mutant strain is able to grow with methanol if the multi-carbon substrate succinate is included in the growth medium as well (22). Under these conditions the *fae* mutant grows at a slower rate and growth arrests at a lower final OD_600_ than the wild-type strain due to accumulation of formaldehyde (22). Kinetic characterization of XoxF1 (this study) and ExaF indicates the capacity of these enzymes to oxidize formaldehyde *in vitro*, and raises the possibility that they may have this activity *in vivo* (5). Should that be the case, one or both of these enzymes could increase the formaldehyde tolerance of the strain when active. We measured growth of the *fae* mutant with La^3+^ (+La) and without La^3+^ (-La) with both succinate and 10 mM methanol as growth substrates [FIG 2B]. Relative to the wild-type strain, the *fae* mutant strain grew more slowly and exhibited a strong reduction in growth rate after surpassing OD_600_ 0.5 in both the +La and -La conditions. After the observed downshift in growth rate, the *fae* mutant strain grew at the same rate regardless of the presence or absence of exogenous La^3+^. However, growth yield, as final OD_600_, was 22% higher in the +La cultures (OD_600_ of 0.8 No La^3+^ vs 1.0 with La^3+^). This is suggestive that *M. extorquens* AM1 is more tolerant of formaldehyde during Ln methylotrophy.

### Impacts of La^3+^ on methanol growth

The effect of Ln on growth rate and yield of methylotrophic bacteria varies by report, ranging from essential for growth (3), to having either a small impact (5, 29), or no impact (2, 18) on methanol growth. Because XoxF1 and MxaFI-catalyzed methanol oxidation produces formaldehyde, it could be expected that Ln-dependent methylotrophy would not differ from methylotrophy without Ln. However, we observed a Ln-dependent increase in formaldehyde tolerance, and since formaldehyde is a key metabolic intermediate in methanol metabolism, we reinvestigated the impact of La^3+^ on methanol growth for the wild-type strain of *M. extorquens* AM1 at both high and low methanol concentrations [Fig 2AC]. All growth media was prepared in new glassware and all cultures in new polypropylene culture tubes to minimize the likelihood of Ln contamination. Using 15 mM methanol as the growth substrate, the wild-type strain grew with specific growth rates of 0.181±0.002 h^−1^ and 0.157±0.003 h^−1^, with (+La) and without (-La) exogenous La^3+^ respectively. This constituted a 15% increase in growth rate with exogenous La^3+^. When the concentration of methanol was increased to 125 mM, the growth rate was 22% faster for the +La condition (μ = 0.180±0.003 h^−1^) relative to the –La condition (μ = 0.146±0.002 h^−1^). Addition of exogenous La^3+^ resulted in a higher culture yield (+12.5%) with 125 mM methanol. A similar La^3+^-dependent increase in yield was observed with 15 mM methanol (+10.0%) and in 50 mL cultures grown in new shake flasks (+17.1%, data not shown). Methanol concentration had no impact on the growth rate for the +La conditions, but increasing to 125 mM methanol resulted in a 7% slower growth rate in the –La condition. Together, these results show that exogenous La^3+^ positively impacts both growth rate and growth yield when methanol is the carbon and energy source. Comparing cell dry weight of methanol grown cultures from shake flasks at full OD, we observed a 21±3% increase (average of 5 biological replicates) in cell mass when exogenous La^3+^ was provided to the medium.

### Methanol, formaldehyde and formate oxidation genes are differentially regulated with Ln

In order to identify changes in methylotrophic metabolism due to the presence of Ln, RNA transcript profiles of cultures grown with and without exogenous La^3+^ were compared using RNA-seq transcriptomics [FIG 3]. In the presence of La^3+^ the genes *xoxF, xoxG*, and *xoxJ* encoding the Ln-dependent MDH, the predicted cognate cytochrome *c*_*L*_, and a MxaJ-like protein, were all significantly upregulated (8-13 fold). Notably, *exaF*, encoding the Ln-dependent ethanol dehydrogenase (5), was upregulated greater than 2-fold suggesting a possible role in Ln-dependent methylotrophy. Expression of *xoxF2* was also upregulated, although the increase in expression was less than 2-fold. In congruence with the “lanthanide-switch” reported for methanol oxidation in methylotrophic bacteria (18, 23, 29), the entire *mxa* gene cluster, encoding the Ca-dependent MDH and accessory genes, was highly downregulated, (388 and 274 fold for *mxaF* and *mxaI* respectively). The genes encoding both *mxcQE* and *mxbDM*, two-component systems necessary for the expression of the *mxa* gene cluster and repression of the *xox1* genes (7), were also downregulated between 2-4 fold. Several genes important for PQQ biosynthesis and processing were downregulated as well, even though XoxF1 and ExaF are PQQ-dependent dehydrogenases. Genes encoding the formaldehyde oxidizing H_4_MPT pathway showed mixed responses. However, in general, the pathway was downregulated.

**FIG 3.**
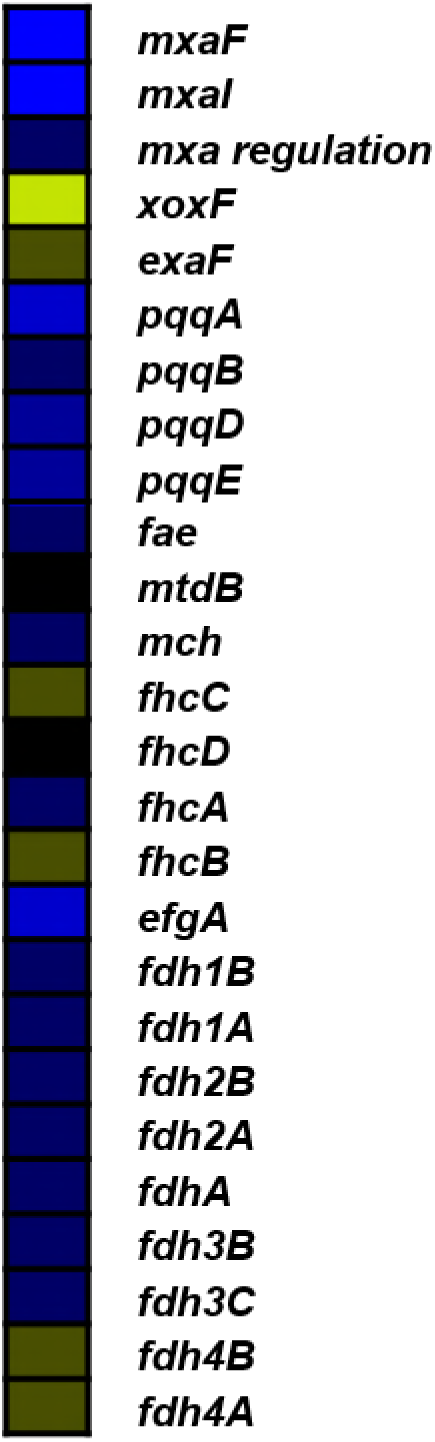
Methylotrophy genes are differentially expressed in the presence of La^3+^. Heat map shows degree of upregulation (yellow) and downregulation (blue) of genes involved in methylotrophic metabolism that were significantly altered in expression in the presence of La^3+^. P-values < 0.05, q-value < 0.1 for all genes shown. Color scale gradations represent log2 fold increments of TMM.

The *M. extorquens* AM1 genome encodes at least four distinct formate dehydrogenases: *fdh1, fdh2, fdh3*, and *fdh4*. The four Fdhs are redundant under standard laboratory conditions, with *fdh4* being the only single knockout that has a measurable impact (30). Fdh3 is predicted to a periplasmic enzyme, and therefore presumed to be necessary for oxidation of formate that would be produced in the periplasm. During La-dependent methylotrophic growth *fdh1, fdh2*, and *fdh3* gene clusters were downregulated 2-3.5 fold, while the *fdh4* gene cluster was upregulated 2-fold, indicating possible changes to formate oxidation. The H_4_F pathway genes showed little change. The genes encoding the assimilatory serine cycle and the EMC pathway exhibited only subtle changes to expression levels. Overall, the gene expression profile of *M. extorquens* AM1 during La-dependent methylotrophy shows clear changes in transcript levels of genes involved in methanol, formaldehyde, and formate oxidation.

### Ln-dependent alterations to methanol, formaldehyde, and formate metabolism

The observed increase in growth rate and yield observed for the wild-type strain in the presence of La^3+^ led us to hypothesize that this could arise from changes in carbon flux to the oxidation pathway (formaldehyde oxidation to formate via the H_4_MPT dependent pathway, where NAD(P)H is produced) vs a two electron transfer from MDH to a terminal oxidase via cytochromes *c*_L_, *c*_H_, and *aa3* complex(31). To assess how methylotrophy is altered in such a way to allow improved growth, we first compared methanol consumption of the wild-type strain in both +La and –La conditions. Wild-type cultures were grown +La and -La with 125 mM methanol to an OD_600_ of 1.0, correlating with mid-exponential growth phase. Growth patterns resembled what we observed with the plate reader for similar conditions. Cells were harvested and residual methanol in the growth medium was quantified by HPLC. The culture medium from the +La culture contained 34% more methanol, corresponding to a 44% reduction in consumption [FIG 4A]. Internal formaldehyde concentrations were not significantly different between the two growth conditions [4B], but we observed a 4-fold increase for the internal formate level for the +La condition [4D]. Excreted formaldehyde and formate concentrations were ∼2-fold higher for the –La condition [4C and 4E]. Together these results show that the presence of La^3+^ in the growth medium results in altered carbon distribution during methylotrophic metabolism, and suggests a bottleneck of carbon flux at formate.

**FIG 4.**
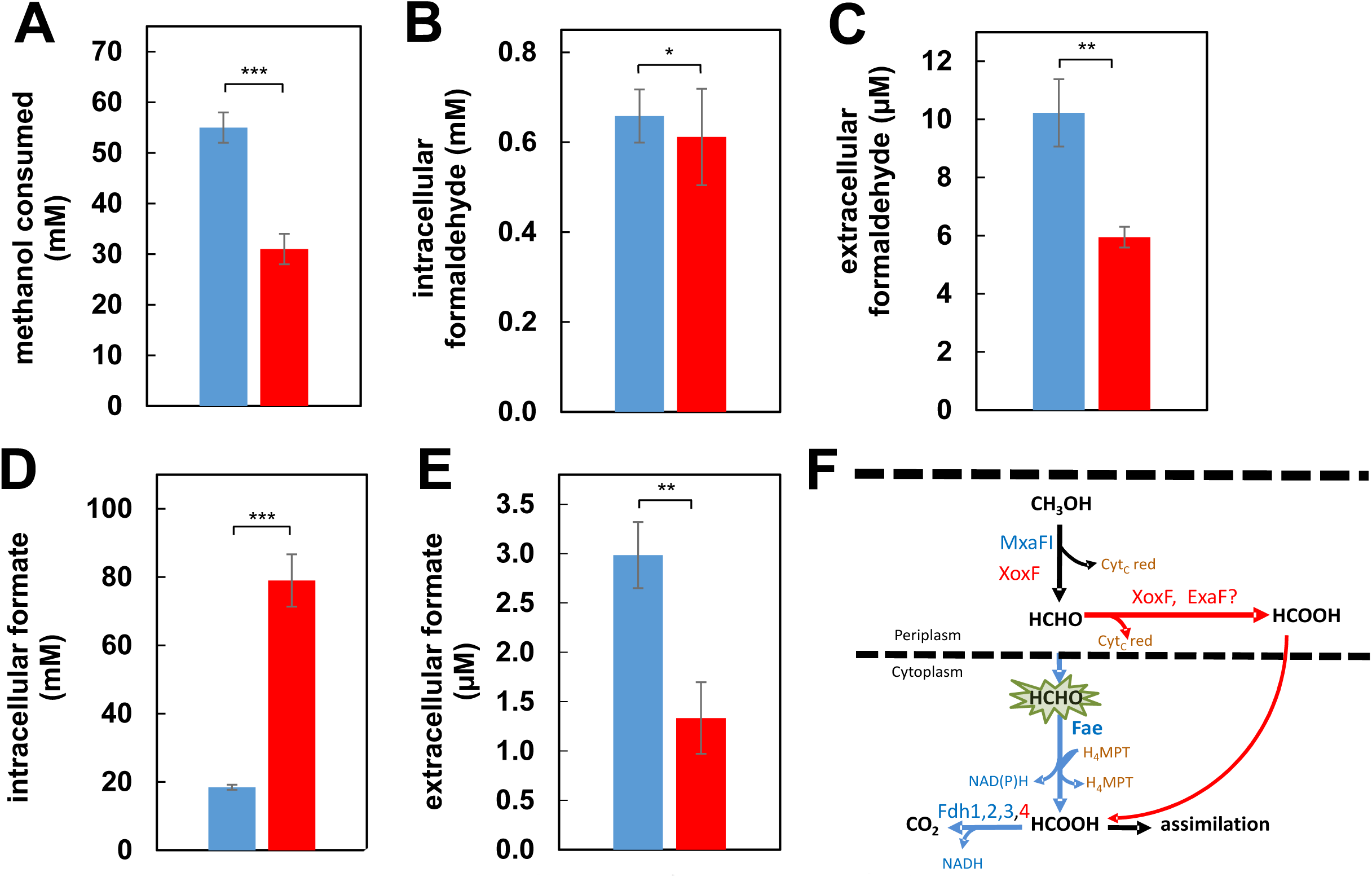
Examination of oxidative metabolism by substrate and product quantification. Cultures were grown in minimal medium with an initial concentration of 125 mM methanol as the growth substrate, and harvested in mid-exponential growth phase at an OD 1.0. The wild-type strain was grown in the absence [blue] or presence [red] of 2 μM exogenous LaCl_3_. (A) Methanol consumption determined by HPLC analysis. Residual methanol in the growth medium at the time of harvest was subtracted from the initial substrate concentration. Values were normalized by subtracting methanol evaporated from uninoculated controls during the same growth period. Values are the mean of 4-6 biological replicates. (B) Intracellular formaldehyde concentrations determined by the Purpald assay. Values represent the mean of three biological replicates. (C) Extracellular formaldehyde concentrations determined from culture supernatants. Values correspond to the average of two biological replicates. (D) Intracellular formate concentrations determined by the NAD^+^-dependent formate dehydrogenase catalyzed formation of a stable formazan. Values are the mean of two biological replicates with three technical replicates each. (E) Extracellular formate concentrations from culture supernatants. Values represent the mean of two biological replicates. (F) Methanol metabolism highlighting methanol to formate oxidation. Blue highlights the enzymes/pathways active during methanol growth in the absence of lanthanides, while red highlights enzymes with increased activity during lanthanide-dependent growth. In all panels, error bars represent the standard deviation for all biological replicates. Multiple-way analysis of variance (ANOVA) was used to determine significance of changes (* = p > 0.05, ** = p < 0.05, *** p = < 0.001).

### Carbon assimilation is altered during Ln-dependent methylotrophy

Next, we assessed how Ln methylotrophy affects the primary assimilatory pathways utilized by *M. extorquens* AM1 for growth on one-carbon substrates using GC-MS and LC-MS/MS targeted metabolomics. Relative pools of key intermediates were compared for cultures grown in the presence and absence of La^3+^ [FIG 5]. Serine cycle metabolite pools were reduced in the +La condition, with the exception of glyoxylate. Reductions in relative pool sizes ranged from 0.4- (serine) to 6-fold (glycerate). Notably, glycerate is phosphorylated to glycerate-2-phosphate, an intermediate that can be siphoned off for C_3_ biosynthesis. Given that glyoxylate is a tightly regulated metabolite that links the serine cycle and EMC pathway, it is not unexpected that the relative pool sizes are similar for both conditions (32, 33). Glyoxylate and acetyl-CoA, the C_2_ intermediate that feeds into the EMC pathway, are produced from malyl-CoA by malyl-CoA lyase. As such, acetyl-CoA pools were relatively unchanged. As with the serine cycle, intermediates of the EMC pathway showed a similar trend with reduced pools in the +La condition for most intermediates. Reductions in EMC pathway intermediate pool sizes ranged from ∼2- to 4-fold. Exceptions to this trend were the mid-EMC pathway intermediates methylsuccinyl-CoA and mesaconyl-CoA, for which the pools were increased ∼2-fold. The TCA cycle intermediate succinate was unchanged, while fumarate and malate were decreased ∼3-fold. Together, these intermediates connect the TCA cycle with the serine cycle and EMC pathway. Notably, citrate levels were increased 2.5-fold, which contrasts with the other organic acid intermediates. Overall, the differences observed show reduced levels of assimilatory intermediates for the two major pathways used to generate biomass precursors during C_1_ growth. It is reasonable then, based on the observed growth and metabolite pool differences, to conclude that these changes reflect more carbon being converted to biomass when La^3+^ is present in the growth medium.

**FIG 5.**
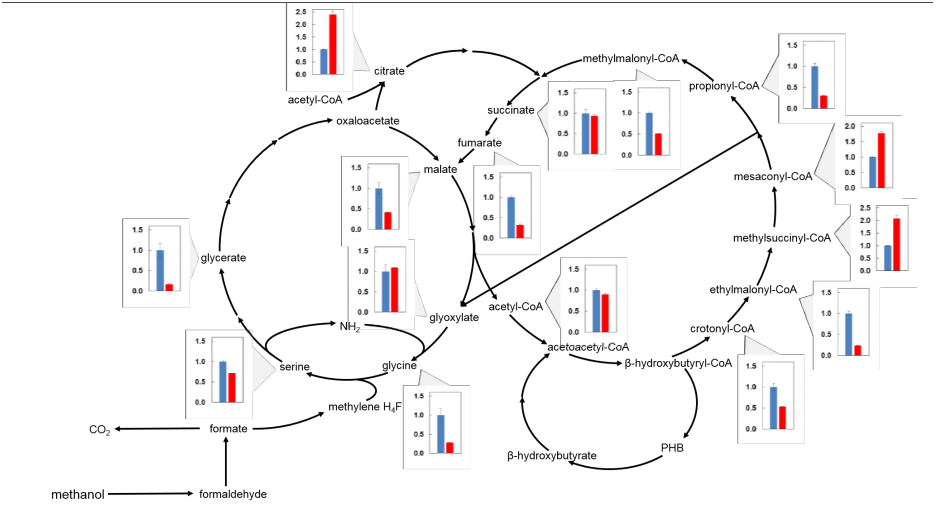
Metabolic intermediates of the assimilatory serine cycle and ethylmalonyl-CoA pathway were measured by mass spectrometry for cultures grown in the absence [-La, blue] or presence [+La, red] of exogenous 2 μM LaCl_3_. Intermediate relative pool sizes are represented as the ratio normalized to the –La condition. Cells were harvested by fast filtration when cultures reached and OD_600_ of ∼0.7-1.1, corresponding with mid-exponential growth phase. CoA intermediates were measured by LC-MS. Organic acid intermediates were measured by GC-MS after TBDMS derivatization. Methysuccinyl-CoA and mesaconyl-Coa, are represented as CoA intermediates in the pathway, but were measured as the derivatized organic acids methylsuccinate and mesaconate, respectively. Error bars represent the SEM of two biological replicates.

### ExaF is a Ln-dependent alternative formaldehyde oxidation system

We observed that a *fae* mutant strain is less sensitive to methanol, and thereby, more tolerant of formaldehyde, when exogenous La^3+^ is available. Our transcriptomics data indicate that during growth with methanol and Ln, *M. extorquens* AM1 produces XoxF1 MDH and the Ln-dependent EtDH ExaF. Although both enzymes in pure form are capable of oxidizing formaldehyde, only ExaF displayed this capability in cell-free extract (this study, (5)). Deletion of *exaF* alone does not impact growth with methanol, with or without Ln (5), but we nonetheless investigated the possibility that ExaF can oxidize formaldehyde during Ln-dependent methylotrophic growth. We conducted the same methanol sensitivity studies with *fae exaF* and *fae* MDH-3 mutant strains (see Table 1) to specifically target the activity of XoxF1 or ExaF, respectively [FIG 6A,B]. The *fae* MDH-3 mutant strain has clean deletions for the *mxaF, xoxF1*, and *xoxF2* genes, but can still produce ExaF. Strains were grown in succinate medium for 2 hours and then 10 mM methanol was added. After addition of methanol, the *fae exaF* mutant strain grew 48% slower and maximum OD was 29% lower than the *fae* mutant strain. Similar effects on growth were seen when the concentration of methanol added was increased to 125 mM, with a 37% reduction in growth rate and 25% reduction in final OD_600_. These results indicate that ExaF is contributing to formaldehyde oxidation *in vivo* in a condition when the intermediate accumulates to toxic levels. They are also further *in vivo* evidence that XoxF1 is not producing formate from the oxidation of methanol. The *fae* MDH-3 mutant strain, on the other hand, displayed a growth pattern resembling the wild-type strain when either 10 mM or 125 mM methanol was spiked into the succinate growth medium [FIG 6A,B], showing that with ExaF available the strain is not inhibited by methanol addition. Measurement of internal formaldehyde concentrations showed that the *fae* mutant strain accumulated 2-fold more formaldehyde compared to the wild-type strain. The *fae exaF* double mutant strain accumulated 26% more formaldehyde than the *fae* mutant strain, indicating that ExaF contributes to oxidize the toxic intermediate *in vivo*. In comparison, >11-fold less formaldehyde accumulated in the *fae* MDH-3 mutant strain compared to the *fae* mutant strain. This is likely the result of ExaF being over-produced in this genetic background, since expression of *exaF* is up-regulated by Ln. The MDH-3 mutant strain also has a severe growth defect with methanol compared to the wild-type, and slower methanol oxidation could contribute as well (18). Overall, these results indicate that ExaF is capable of *in vivo* formaldehyde oxidation during Ln methylotrophy, possibly functioning as an auxiliary oxidation system to prevent inhibitory or lethal accumulation of this toxic intermediate.

**TABLE 1:** Strains and plasmids used in this work

| Strain of plasmid | Description | Reference |
| --- | --- | --- |
| Strains |  |  |
| *Escherichia coli* |  |  |
| DH5α | electrocompetent cloning strain | Invitrogen |
| S17-1 | conjugating donor strain | (55) |
| *Methylobacterium extorquens* |  |  |
| AM1 | wild type; rifamycin-resistant derivative | (56) |
| *exaF* | $\Delta exaF$ | this work |
| *fae* | $\Delta fae$ | (22) |
| *fae exaF* | $\Delta fae \Delta exaF$ | this work |
| *fae* MDH-3 | $\Delta fae \Delta mxaF \Delta xoxF1 \Delta xoxF2$ | this work |
| MDH-3 | $\Delta mxaF \Delta xoxF1 \Delta xoxF2$ | (18) |
| ADH-4 | $\Delta mxaF \Delta xoxF1 \Delta xoxF2 \Delta exaF$ | this work |
| Plasmids |  |  |
| pCM157 | *cre* expression plasmid, Tc<sup>R</sup> | (57) |
| pRK2013 | helper plasmid, IncP *tra* functions, Km<sup>R</sup> | (58) |
| pLB01 | Km<sup>r</sup>, *P<sub>xox1</sub>-xoxF1*-hexahistidine tag, Xa site | (5) |
| pNG284 | pLB01 with TEV cleavage site | this work |
| pHV24 | pCM184:*exaF*; donor for *exaF*::Km | (5) |
| pHV2 | sacB-based allelic exchange *fae* donor; Km<sup>R</sup> | E. Skovran |

**FIG 6.**
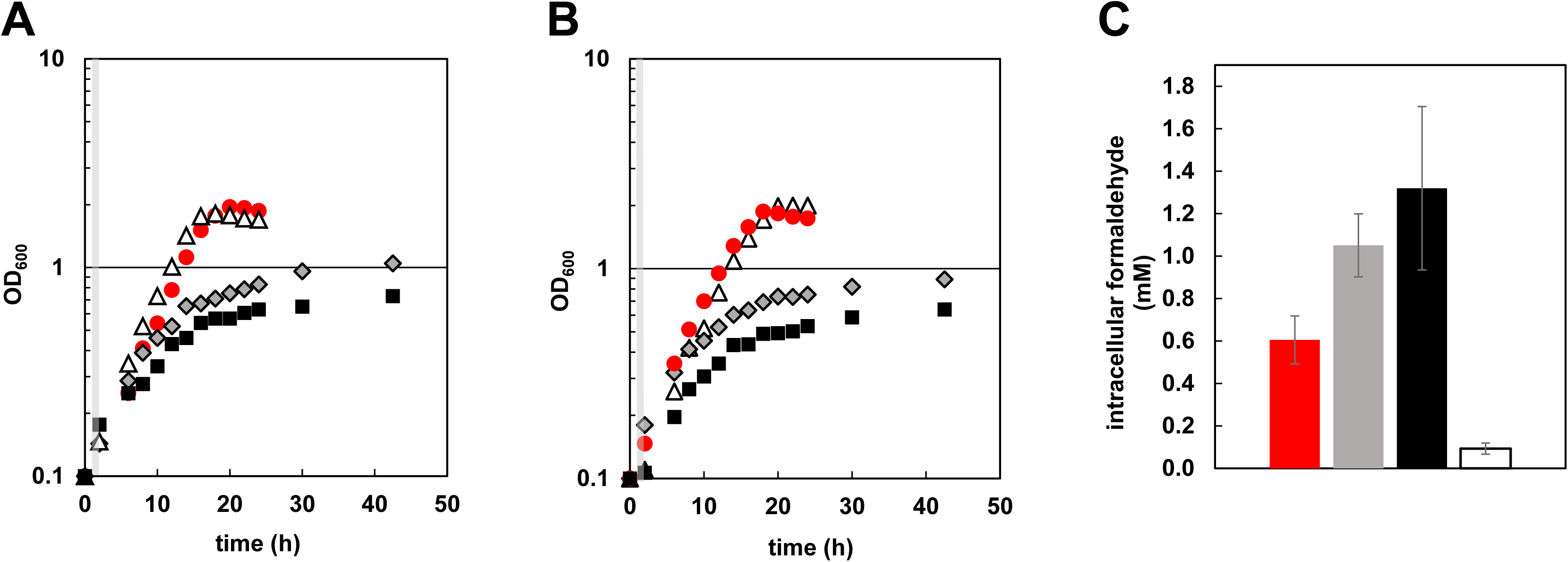
Growth of wild-type and mutant strains on succinate with methanol addition. OD at 600 nm was recorded over time to monitor growth and is reported in log scale. Time points represent three biological replicates with variances <5%. All strains were grown in plastic 15 mL culture tubes. (A and B) Wild-type [red circles], Δ*fae* [gray diamonds], Δ*fae* MDH-3 [white triangles], and Δ*fae* Δ*exaF* [black squares] mutant strains were grown in the presence of 2 μM LaCl_3_ and methanol (A, 10 mM or B, 125 mM) added after two hours of growth with succinate (gray bars). (C) Internal formaldehyde concentrations, determined by Purpald, for wild-type (red), *fae* (gray), *fae exaF* (black), and *fae* MDH-3 mutant strains. Values represent the average of 3-5 biological replicates with standard deviations shown (error bars). Samples were collected at optical densities corresponding to “mid-exponential” growth depending on the strain. Collection ODs were: wild-type, 1.0; *fae* 0.4; *fae exaF*, 0.4; *fae* MDH-3, 1.0.

## DISCUSSION

We have shown that XoxF1-catalyzed oxidation of methanol in *M. extorquens* AM1 produces formaldehyde, distinguishing the Type V XoxF MDH from the Type II XoxF MDH from *M. fumariolicum* SolV (3), and demonstrating that not all XoxF MDH produce formate as the final product of methanol oxidation. Kinetic characterization in this study reveals a 70 to 90-fold increase in XoxF1 catalytic efficiency with La^3+^or Nd^3+^ as a cofactor, relative to no Ln cofactor. The exceptional catalytic efficiency observed for the Type II MDH from *M. fumariolicum* SolV is the result of very high affinity of methanol as the substrate (3). Intriguingly, the higher efficiency for XoxF1 in *M. extorquens* AM1 is instead attributable to increased enzyme activity. In fact, the *K*_M_ for methanol increased 3 to 4-fold with a Ln cofactor. The *K*_M_ for methanol of XoxF1-Nd is ∼2-fold lower than that of XoxF1-La, suggesting that if the enzyme were fully loaded with Nd^3+^ the efficiency would be higher. Further studies with various Ln cofactors are needed to fully understand how these metals impact kinetic properties of XoxF MDH and Ln ADH in general. Detailed structural and/or biochemical studies may be necessary to pinpoint the features or mechanisms responsible for differing oxidation end products among XoxF MDH families. Nonetheless, the current study provides novel insight to our limited understanding of the impact of Ln on catalytic properties of PQQ ADH and highlight the necessity for biochemical studies of representatives from all five families of XoxF MDH.

Using a multi-layered approach, we showed that Ln-dependent methylotrophy in *M. extorquens* AM1 involves significant changes in oxidative and assimilatory metabolism, and results in more efficient growth and higher biomass. Oxidation of methanol by *M. extorquens* AM1 produces formaldehyde, and either through the H_4_MPT pathway and/or ExaF, formate. Production and consumption of these intermediates is intricately balanced and potentially impacts energy conservation and the generation of reducing power as NAD(P)H (34). Periplasmic production of formate from methanol would result in the loss of formaldehyde, and as a result a reduction of flux through the direct and primary route for production of NAD(P)H as the product of methylene tetrahydrofolate (H_4_F) dehydrogenases, MtdA and/or MtdB [Fig 4F]. NADPH is known to drive assimilation and define growth rate (35). It is not surprising that the H_4_MPT pathway is indispensable even if periplasmic formate production is possible, and indeed we observed no differences in the internal formaldehyde pools of wild-type cultures grown with and without exogenous La^3+^.

The increase of internal formate in the +La cultures suggests a redistribution of carbon at this important metabolic branchpoint. Several factors could contribute to the accumulation of formate during Ln-methylotrophy. It is possible that XoxF1 is a more efficient enzyme than MxaFI, thereby more readily providing formaldehyde as substrate for Fae. Indeed we observed an increase *V*_max_ and catalytic efficiency for XoxF1 with Ln. We also reported a significant increase in MDH activity from wild-type cell-free extracts grown with La^3+^ [p < 0.01, one-way ANOVA] (18). We have also shown evidence that XoxF and ExaF are capable of producing formate from formaldehyde, which could provide a H_4_MPT-independent route for formate production. Serine hydroxymethyl transferase (GlyA) has been identified as a potential control point for growth with methanol based on relatively large pool sizes of glyoxylate and glycine substrates and enzyme activity below the theoretical minimum activity needed to support growth (36, 37). It is likely that GlyA serves as a metabolic control point during Ln-methylotrophy as well. The H_4_F reactions are reversible, while the H_4_MPT reactions are not, providing three possible fates for formate. We observed that excreted formate was reduced 2-fold in the +La cultures. It is possible that formate oxidation to CO_2_ is reduced. During canonical methylotrophy, carbon flux through formate is split with ∼70% being oxidized to CO_2_ and ∼30% going to assimilation (28). The observed down-regulation of Fdh1, Fdh2, and Fdh3-encoding genes, and up-regulation of the genes encoding Fdh4, is suggestive of a regulatory switch at this metabolic branchpoint that may drive more efficient assimilation of carbon and correlate to the observed higher biomass and growth rate. Changes in NADH/NAD ratios are not known to impact growth rate in *M. extorquens* AM1 (35), and Fdh4 is the only formate dehydrogenase that is essential for normal methanol growth (30).

Targeted metabolomics analysis of the assimilatory pathways revealed decreased pools for several serine cycle and EMC pathway intermediates. Interestingly, the EMC pathway intermediates methylsuccinyl-CoA and mesaconyl-CoA showed increased pools. The latter intermediate is converted to β-methylmalyl-CoA by mesaconyl-CoA dehydratase, which was identified as a potential carbon flux control point during a transition from succinate growth to methanol growth (38). Larger pools of mesaconyl-CoA and methylsuccinyl-CoA, suggest Mcd may function as a control point for EMC pathway flux during Ln-dependent methylotrophy. Confirmation of the possible control points mentioned above would require re-examining enzyme activities and/or fluxes for the serine cycle and EMC pathway and is beyond the scope of this study. An additional observation from our metabolomics is the 2.5-fold relative increase of citrate that can be excreted to chelate trace metals needed for metabolism (39). As the mechanism for Ln acquisition is still unknown, excretion of citrate to solubilize Ln for uptake is formally possible, and a potential candidate compound for future investigation. Overall, our metabolomics assessment of the assimilatory pathways shows a distinctive pattern for Ln-dependent methylotrophy that is consistent with the growth phenotypes observed.

*Methylobacterium* strains are associated with the plant phyllosphere, a dynamic environment where competition for volatile organic substrates, such as methanol and ethanol, is expected to be high (40–42). We show that multiple ADH systems and regulatory mechanisms could function together to maintain stability of metabolism in an environment with large fluctuations in substrate availability. Such complementarity of oxidation systems may further explain the large number of bacteria replete with redundant enzymatic systems. It has been speculated that redundant, cofactor-dependent oxidation systems may allow for effective growth under differing metal limitation. We observed up-regulation of two independent Ln ADH systems, XoxF1 MDH and ExaF EtDH in the presence of Ln, and a significant positive impact of Ln on bacterial metabolism. Ln, therefore, may cause a shift in metabolic strategy to one that is more efficient. The underlying metabolic tradeoffs of such a switch are still unknown.

However, expression of PQQ synthesis genes was down-regulated in the presence of Ln. This is in agreement with the recent study comparing gene expression patterns in *M. aquaticum* strain 22A (19). Why PQQ synthesis is down-regulated with the up-regulation of PQQ ADHs in the presence of Ln is unknown. Activities of XoxF1 and ExaF do not appear to be impacted by the potential down-regulation of PQQ synthesis, suggesting that any decrease in PQQ pools is not severe enough to restrict formation of holoenzyme. PQQ is a known plant growth promotion factor (43), however, and it is possible that restricting PQQ plays a regulatory role, either with other enzymes of the microbe, the plant, or both. Further studies of PQQ in Ln methylotrophy are needed to determine the regulatory role of this molecule.

## MATERIALS AND METHODS

### Bacterial strains and cultivation

Strains and plasmids used in this study are listed in TABLE 1. *Escherichia coli* strains were maintained on solidified (1.5% wt/vol agar) Luria-Bertani (LB) medium (44) (BD, Franklin Lakes, NJ) at 37 °C with kanamycin added to a final concentration of 50 μg/mL. *M. extorquens* AM1 strains were grown in minimal salts medium (45) prepared in new glassware that had not been exposed to Ln. Precultures were grown in round-bottom polypropylene culture tubes (Fisher Scientific, Waltham, MA, USA) at 30 °C, shaking at 200 rpm on an Innova 2300 platform shaker (Eppendorf, Hamburg, Germany), with succinate (15 mM) or methanol (125 mM) as the growth substrate. LaCl_3_ was added to a final concentration of 2 μM or 20 μM when indicated. When necessary, kanamycin was added to a final concentration of 50 μg/mL. For methanol tolerance studies, methanol was added to a concentration of 10 mM or 125 mM.

### Plasmid and strain construction

Plasmid pLB01 was constructed for producing hexahistidine-tagged XoxF1 with a Factor Xa protease cleavage site (5). For this work, the Factor Xa cleavage site in pLB01 was substituted with the Tobacco Etch Virus protease cleavage site (25) to generate pNG284 using the following primers: pLB01 vector backbone; forward GCCGAACAACGGATTGGAAGTACAGGTTCTCCATCATCACCATCACCATAATTGTC, reverse CAGCTCACTCAAAGGCGGTAATAC; *xoxF1* allele insert; forward, CGTATTACCGCCTTTGAGTGAGCTGCTGAATTTAGCAGGCAAGTTTCCTG, reverse, GATGGAGAACCTGTACTTCCAATCCGTTGTTCGGCAGCGAGAAGAC. Primers were designed to generate PCR products with 20 bp homologous overlapping regions between backbone and insert fragments. The backbone forward primer and insert reverse primer were designed to contain the TEV protease cleavage site. Vector backbone and allele insert PCR products were assembled using *in vivo* gap repair assembly (46, 47). Briefly, *E. coli* DH5α electrocompetent cells were transformed via electroporation with ∼150 ng of backbone and insert PCR products. Transformed cells were incubated at 37 °C for ∼1 h outgrowth and then plated on LB plates with kanamycin to select for transformants. Deletion mutant strains were constructed as reported (5).

### Phenotypic analyses

To compare growth among strains, cultures were grown overnight in succinate minimal media. Overnight cultures were centrifuged at 10,000 × *g* at room temperature for 2 minutes. Spent culture medium was removed, cells were washed twice with fresh carbon-free culture medium, and then resuspended in fresh carbon-free culture medium. Growth phenotypes of wild-type *M. extorquens* AM1 were compared using a BioTek EpochII microplate reader (BioTek, Winooski, VT) following the procedures described (45) with slight modifications. Briefly, 650 μL of growth medium with methanol, with or without 2 μM LaCl_3_, was inoculated to a starting optical density (OD) of 0.1. Cultures were shaken at 548 rpm at 30 °C and the OD at 600 nm was monitored over time. OD measurements were fitted to an exponential model for microbial growth using CurveFitter (http://www.evolvedmicrobe.com/CurveFitter/). Growth curves were performed in quadruplicate for each condition.

*M. extorquens* strains were tested as previously described for sensitivity of methanol (22). Precultures were grown overnight with succinate in polypropylene tubes. Three-mL cultures of minimal medium with succinate in polypropylene tubes were inoculated from precultures to a starting OD_600_ of 0.1 and shaken continuously at 200 rpm and 30 °C. After 2 h, methanol or water was added as a 100 μL volume to the culture tubes to final concentrations of 0 mM, 10 mM or 125 mM methanol. After additional of methanol, cultures were continuously shaken and growth was monitored by measuring OD_600_ every 2 h for 24 h using an Ultraspec 10 cell density meter (Amersham Biosciences, Little Chalfont, UK). Methanol inhibition studies were performed in triplicate for each condition.

### RNA-seq transcriptomics

50 mL cultures were grown in shake flasks with or without LaCl_3_ to an OD_600_ of 0.7 that correlated with mid-exponential growth. Total RNA samples were procured, and quality was verified as previously described (48). Two biological replicates were prepared for each condition. rRNA depletion, library preparation, and Illumina Hi-Seq sequencing were performed by the Michigan State University Research Technology Support Facility Genomics Core. Libraries were prepared using the TruSeq Stranded Total RNA kit (Illumina, San Diego, CA), with Ribo-Zero Bacteria used for rRNA depletion (Epicentre, Madison, WI). All replicates were sequenced on an Illumina HiSeq 2500 using a multiplex strategy with 50 bp single-end reads with a target depth of >30 million mapped reads. Base calling was done by Illumina Real Time Analysis (RTA) v1.18.64 and the output of RTA was demultiplexed and converted to a FastQ format with Illumina Bcl2fastq v1.8.4. The filtered data were processed using SPARTA: Simple Program for Automated for reference-based bacterial RNA-seq Transcriptome Analysis (49). Final abundances were measured in trimmed mean of M values (TMM).

### Methanol consumption measurements

Shake flasks were cleaned of potential Ln contamination by repeatedly growing an *mxaF* mutant until the strain no longer grew (18). Then, flasks containing 50 mL minimal medium with 125 mM methanol were inoculated with succinate-grown precultures. Cultures were grown shaking continuously at 200 rpm and 30 °C to OD_600_ of 1.0, corresponding to mid-exponential growth. The culture was then transferred to a 50 mL conical tube (Fisher Scientific, Waltham, MA, USA) and cells were pelleted using a Sorvall Legend X1R centrifuge (Thermo Fisher Scientific, Waltham, MA) at 4,696 × *g* at 4 °C for 10 minutes. One mL of supernatant was centrifuged at room temperature for 25 minutes at 15,500 × *g*, and then transferred to a clean glass HPLC vial for immediate analysis. Samples were analyzed using a Shimadzu Prominence 20A series high-pressure liquid chromatography system with an SPD-20A UV-VIS detector (Shimadzu, Kyoto, Japan) and a BioRad Aminex HPX-87H organic acids column 300 × 7.8 mm with a 9 µm particle size (BioRad, Hercules, California, USA). An isocratic flow of 5 mM HPLC grade sulfuric acid in Nanopure water was used as the mobile phase at 0.6 mL/min. Peak areas were compared with a standard curve to determine methanol remaining in the media. Due to the volatility of methanol, [methanol consumption] values were normalized by subtracting methanol concentrations of samples taken from uninoculated control flasks at the same time

### Intracellular formaldehyde and formate measurements

To determine internal concentrations of formaldehyde, cells were resuspended in 2 mL 25 mM Tris-HCl pH 8.0 with 150 mM NaCl and broken using a One Shot Cell Disruptor set to 25 PSI (Constant Systems, Ltd., Daventry, UK). Cell lysates were centrifuged at 15,500 rpm for 5 min at 4 °C. The supernatant was transferred to a clean 1.5 mL tube and the formaldehyde concentration was determined using the Purpald assay (50) and measuring absorbance at 550 nm using a BioTek EpochII microplate reader (BioTek, Winooski, Vermont, USA). Internal formate concentrations were measured by the formate dehydrogenase catalyzed conversion of formate to a stable formazan color (51). Briefly, cells were resuspended and lysed in 2 mL 50 mM phosphate buffer, pH 7.4. After centrifugation to remove cell debris, 50 μL of the sample was added to a reaction mixture containing 100 μL 50 mM phosphate buffer, pH 7.4, 0.04 U NAD^+^-dependent formate dehydrogenase from *Candida boidinii* (MilliporeSigma, St. Louis, MO USA), and 100 μL of color reagent containing 1.9 mmol iodonitrotetrazolium (INT), 1.2 mmol β-NAD^+^, and 0.065 mmol phenazine-methosulfate (PMS). The reaction mixture was incubated at 37 °C for 30 minutes, quenched with 0.1 M HCl, and absorbance was measured at 510 nm. For internal concentration estimation, a dry weight of 0.278 g/L at 1 OD_600_ unit was used, as previously reported (28), with a cell volume of 36 μL/mg dry weight, based on an average cell size of 1 by 3.2 μm (52) and an average of 4×10^8^ cells/mL at 1 OD_600_ unit (53).

### LC-MS/MS and GC/MS Metabolomics

To measure the accumulation of serine cycle and EMC pathway intermediates, samples were prepared from 30 mL of culture harvested via vacuum fast filtration during mid-exponential growth phase (OD_600_ = 0.7 − 1.1), transferred to a 50 mL conical polypropylene tube, immediately frozen in liquid N_2_, and stored at −80 °C until use. Hot extraction using 30 mL of boiling water was achieved as described before (54). Internal standards were added by pipetting directly to each filter before extraction, 10 μL of 25 μM acetyl-1,2-^13^C_2_-CoA and 15 μL of 300 μM Cell Free Amino Acid Mixture (MilliporeSigma, St. Louis, MO) containing L-Threonine-^13^C_4_,^15^N. Dried samples were resuspended in 70 μL of ddH_2_O, and 30 μL was used for CoA derivative analysis. From the remaining sample, 30 μL were concentrated in the speed vac to dryness and used for organic acid derivatization. Methoximation was performed overnight at 37 °C after addition of 20 μL of 20 mg/mL O-methyoxamine HCl in pyridine. *Tert*-butyldimethylsilyl trifluoromethanesulfonate (TBMDS) silylation was achieved by addition of 20 μL MTBSTFA incubation at 60 °C for 1h. Derivatized organic acids were measured with an Agilent 5975 GC single quadrupole mass spectrometer (Agilent Technologies, Santa Clara, California, USA) using a 30-m VF5-ms column. CoA derivatives were measured by LC-MS/MS using Waters Xevo TQD tandem quadrupole mass spectrometer (Waters Corporation, Milford, Massachusetts, USA) and an Acquity UPLC CSH C18 column (2.1×100 mm) with a 1.7 µm pore size (Waters Corporation, Milford, Massachusetts, USA). Aqueous Mobile Phase A was 400 mM ammonium acetate in water with 1% formic acid and organic Mobile Phase B was acetonitrile using a flow rate of 0.4 mL/min, with the following gradient: 0-1 min 98% aqueous phase, 1-3 min gradient to 95% aqueous phase, 3-5 min gradient to 60% aqueous phase, 5-6 min maintain flow at 40% aqueous phase, 6-8 min maintain flow at 98% aqueous phase. LC-MS/MS dwell times were optimized so all compounds were within the instrument’s range of detection and multiple reaction monitoring (MRM) channels were as follows: acetyl-CoA 810.1>303.1; acetyl-1,2-^13^C_2_-CoA 812.1>305.1; propionyl-CoA 824.1>317.1; crotonyl-CoA 836.1>329.1; methylmalonyl-CoA 868.1>317.1; ethylmalonyl-CoA 882.2>375.2. All data were analyzed with QuanLynx, where ions corresponding to loss of the *tert*-butyl group [M-57]^+^, and *m/z* = 160 for methoximated glyoxylic acid (1TBMDS, 1MO) were used during GC-MS data analysis.

### Protein purification and analysis

2.8-L shake flasks cleaned for Ln contamination as described above were used. Cultures of wild-type *M. extorquens* AM1 harboring pNG284 were grown with methanol and 20 μM LaCl_3_ to OD ∼6. XoxF1 was purified by IMAC and processed, including determination of metal content by inductively-coupled plasma mass spectrometry (ICP-MS) and PQQ content by UV-visible spectroscopy, as reported in (5).

### Methanol dehydrogenase activity assay and enzyme kinetics

Methanol dehydrogenase activity was measured using the PMS reduction of 2,6-dichlorophenol-indophenol (DCPIP) according to Anthony and Zatman, with modifications reported in Vu *et al.* (10, 18). Using these modifications, there was little to no endogenous reduction of DCPIP without addition of methanol. The assay parameters, therefore, were suitable for determining kinetic constants for XoxF1. Kinetic constants were determined by measuring enzyme activity with a range of substrate concentrations. Data were fitted and constants calculated according to the Michaelis-Menten equation using Prism GraphPad 6 (GraphPad Software, La Jolla, CA).

## ACKNOWLEDGMENTS

We would like to thank E. Skovran for providing the pHV2 *fae* allelic exchange plasmid. We thank Michaela TerAvest and Paula Roszczenko for critical revision of this manuscript. Mass spectrometry analyses were conducted in the Michigan State University Mass Spectrometry and Metabolomics Core Facility with special thanks to Scott Smith for instrument and analytical training. Metal content measurements were performed at the Michigan State University Laser Ablation ICP-MS Facility.

## FUNDING SOURCES

This material is based upon work supported by the National Science Foundation under Grant No. 1750003.

---

**FIG S1** Dataset of transcriptomics analysis. All replicates were sequenced on an Illumina HiSeq 2500 using a multiplex strategy with 50 bp single-end reads with a target depth of >30 million mapped reads. Base calling was done by Illumina Real Time Analysis (RTA) v1.18.64 and the output of RTA was demultiplexed and converted to a FastQ format with Illumina Bcl2fastq v1.8.4. The filtered data were processed using SPARTA: Simple Program for Automated for reference-based bacterial RNA-seq Transcriptome Analysis (49). Final abundances were measured in trimmed mean of M values (TMM). No La represent data sets from cultures grown with no lanthanum and methanol; Plus La represent data sets from cultures grown with methanol and lanthanum; BR numbers refer to biological replicates; logFC is the log fold change between the two conditions; logCPM are the values representing the log counts per million reads; LR represents the log ratio; and FDR is false discovery rate.

**FIG S2** SDS-PAGE gel analysis of XoxF1 MDH with Ln cofactors. 1.5 L cultures of wild-type *M. extorquens* AM1 harboring pNG284 were grown with methanol in 2.8 L shake flasks cleaned of residual Ln by growth of an *mxaF* deletion mutant. Cleaning of glassware was repeated until the *mxaF* mutant did not grow for > 48 h. After inoculation of the wild-type/pNG284 strain, 20 μM LaCl_3_ or NdCl_3_ was added to the growth medium and cultures were harvested at an OD ∼6. XoxF1 was purified by IMAC using Ni-NTA Superflow resin (Qiagen, Hilden, Germany). The hexa-histidine affinity tag was removed using TEV protease, and protein purity and size was analyzed by SDS-PAGE.

